# From nodes to pathways: an edge-centric model of brain function-structure coupling via constrained Laplacians

**DOI:** 10.1101/2024.03.03.583186

**Authors:** Viljami Sairanen

## Abstract

Understanding how functional relationships relate to the brain’s structural architecture remains a central challenge in network neuroscience. Many Laplacian-based approaches describe function-structure coupling at the node level, which can make it difficult to identify the specific anatomical pathways that support observed functional relationships. This work introduces a constrained Laplacian formulation that incorporates externally specified pairwise functional relationships and yields a nodal field whose graph-gradient representation produces edge-level quantities describing how the structural network accommodates these relationships. The method is implemented using Modified Nodal Analysis, enabling efficient computation on large connectomes. Given an observed pattern of functional associations and a structural connectivity graph, the proposed framework estimates which structural edges are most consistent with supporting the imposed pattern. The framework is demonstrated in multiple settings, including a single-subject example, a controlled diffusion phantom, an in-silico function-structure simulation, and analyses of Human Connectome Project data at both group level (207 subjects) and test-retest conditions (three subjects). Across these applications, the method produces an edge-level representation of function-structure coupling, enabling pathway-specific analysis of brain connectivity.

## 1 Introduction

Understanding how structural connectivity (SC) relates to functional connectivity (FC) is a central challenge in network neuroscience (Greaves et al., 2025). SC provides the anatomical scaffold of axonal pathways, often measured using diffusion-weighted MRI tractography, whereas FC is typically characterized as statistical dependencies in neural activity measured via fMRI, EEG, or MEG. Although SC and FC exhibit macroscale correlations (Honey et al., 2009), many functional relationships occur without a direct structural link, reflecting polysynaptic and network-mediated interactions and motivating computational models that formalize this coupling.

A prominent class of approaches to couple SC and FC relies on spectral graph theory (Chung, 1997), which uses the graph Laplacian of the SC network and its eigendecomposition to model functional relations. For example, network diffusion models (Abdelnour et al., 2014) predict FC by simulating heat-like diffusion on the SC graph, while spectral models (Abdelnour et al., 2018; Atasoy et al., 2016; Deslauriers-Gauthier et al., 2020) express FC patterns as combinations of connectome harmonics. These harmonics correspond to Laplacian eigenmodes that capture indirect pathways and polysynaptic interactions. These approaches exploit the eigendecomposition of the Laplacian to reveal fundamental modes of network communication, providing a principled link between structure and function grounded in graph signal processing (Shuman et al., 2013).

Despite their success, existing Laplacian related network models primarily yield node level descriptions of function-structure coupling, in which functional interactions are summarized between pairs of regions. While such models implicitly account for multi-edge pathways, they do not explicitly identify which structural connections or tractography streamlines support a given functional relationship. As a result, the contributions of individual edges or specific structural pathways to functional correlations remain latent and difficult to interpret. Moving beyond node-level representations toward an edge-centric perspective offers a route-based view of function-structure coupling, enabling the tracing of structural support patterns through concrete anatomical pathways.

In this work, the term “node-level” is used to refer to connectivity measures defined between pairs of regions, such as conventional functional connectivity matrices where each value summarizes the relationship between two brain areas. In contrast, “edge-level” outputs refer to quantities defined on individual structural connections or tracts. For example, a node-level analysis might report the functional connectivity between the posterior cingulate cortex and medial prefrontal cortex, whereas an edge-level analysis can describe which specific white-matter pathways between these regions contribute most strongly to that relationship.

A wide range of alternative computational approaches have also been proposed to link brain structure and function. Some model biophysical dynamics (Breakspear et al., 2010; Deco, 2015; Liu et al., 2022), including the generation of realistic neural time-series, while others perform statistical analyses of relationships between structural and functional connectomes (Huang and Ding, 2016; Suárez et al., 2020), or apply machine learning methods such as Graph Neural Networks (GNNs) (Chen et al., 2024; Hong et al., 2023; Neudorf et al., 2022; Yang et al., 2022; Zalesky et al., 2024). These approaches differ in their formulation and in the level at which function-structure relationships are modeled, ranging from time-series dynamics to statistical dependencies and derived connectivity measures. However, the applications that summarize interactions at the level of region pairs, tend to have limited capability of identification of the specific structural pathways supporting a given functional relationship.

Complementary lines of work have also projected functional information onto structural pathways. Track-weighted functional connectivity (Calamante et al., 2013) combines voxel-wise FC maps with tractography streamlines to highlight white-matter segments associated with specific networks. Deslauriers-Gauthier et al. (2019) infer time-resolved, directed interactions by integrating diffusion MRI with EEG to model information flow supported by structural bundles. Spectral Dynamic Causal Modeling (Novelli et al., 2024) instead fits a state-space generative model to cross-spectral features of fMRI, providing estimates of directed effective connectivity. These approaches offer voxel-wise, dynamic, or directed perspectives on function-structure relationships. The present framework differs by formulating SC-FC coupling as a constrained Laplacian problem, yielding a pathway-resolved representation directly tied to the network’s edge structure while remaining analytically transparent and connectome-wide.

This study builds on graph-Laplacian approaches and introduces a framework that moves beyond node-level representations to characterize how functional relationships are supported across the edges of the structural connectome. By formulating function-structure coupling as a constrained Laplacian problem, functional relationships are mapped onto structural pathways, yielding interpretable measures of how these relationships are supported across the structural network. The proposed edge-centric perspective adds a new dimension to graph-based network analyses by providing connection and tractography streamline specific insights that may help in uncovering mechanisms underlying brain disorders and conditions. Beyond its theoretical contribution, the framework enables practical applications such as identifying structural connections that preferentially support functional coupling and informing tractography filtering using functional data. To facilitate adoption, an open-source Python implementation for applying this framework to existing SC and FC datasets is freely available in Github^1^.

## 2 Theory

Spectral graph theory provides a principled framework for modeling signal interactions within networks, with the graph Laplacian encoding conservation of flow across nodes (Chung, 1997; Shuman et al., 2013). In network neuroscience, Laplacian eigendecomposition has been used to characterize function-structure relationships at the graph node level (Honey et al., 2009).

This section builds on this theory and describes an idea of how the Laplacian eigen-decomposition could be extended to define nodal potentials originating from the functional brain signals whose graph gradients result in edge-level patterns describing how functional information is distributed within the structural network.

### 2.1 Graph Laplacian equality between node and edge spaces

The structural connectome can be represented as an undirected weighted graph *G* = (*V, E, W*) where *V* denotes the set of vertices corresponding to brain regions in the parcellation, *E* are edges i.e. white-matter connections between the regions, and *W* is the weighted adjacency matrix that encode nodewise structural connection strength such as number of streamlines, for example, and is given by:

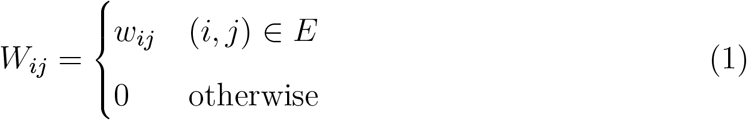

The degree matrix of graph *G* is given by *D* = diag(∑_*j*_ *w*_*ij*_). Throughout this work, the graph *G* is undirected and the Laplacian is symmetric. Accordingly, the formulation does not infer directed or source-to-target influences.

In the case of weighted graph, the graph Laplacian operator acts on nodewise signals as (ℒ *f*) (*i*) = _*j*_ *w*_*ij*_ (*f* (*i*) − *f* (*j*)) and leads to two equivalent matrix representations of the graph Laplacian. The first representation in node space:

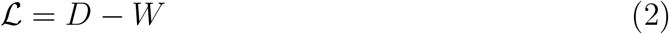

and the second representation is based on incidence-based factorization:

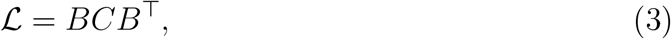

where *B* is oriented incidence matrix and *C* = diag(*e*_*w*_), *e* ∈ *E* is edge weight matrix (Chung, 1997; Shuman et al., 2013). Structural edge weights *C* quantify how strongly each connection contributes to differences between neighboring nodes. The expression *BCB*^⊤^ is a structural decomposition that exposes how node-level differences are shaped by the network’s edges giving equality

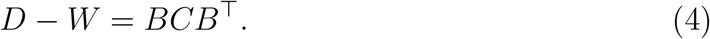

### 2.2 From Laplacian eigenmodes to nodal potentials

The equivalence in eq. 4 shows that the graph Laplacian admits complementary node-based and edge-informed representations. In the node-space formulation, the Laplacian characterizes how signals defined on regions differ from their neighbors, whereas in the edge-informed formulation it encodes how such differences are distributed along individual structural connections.

To relate graph Laplacian operator to brain functions, consider a scalar-valued nodal field *ϕ* ∈ ℝ ^|*V* |^. This can be interpreted as a nodal potential originating from the functional information, and it can be solved from the following Laplacian system

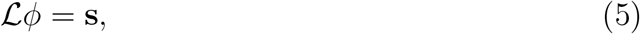

where **s** ∈ ℝ ^|*V* |^ represents nodal source terms that encode functional constraints or external inputs imposed on the structural network. This formulation does not model the propagation or inference of functional signals, nor does it attempt to estimate or reproduce functional connectivity. Instead, it specifies a forward constrained Laplacian problem in which selected functional connectivity values act as externally imposed relationships. The unknowns in this system are the nodal potentials *ϕ* and the resulting edge flows, while the structural Laplacian ℒ and the functional constraints **s** are observed inputs. The solution therefore represents how the structural network would support the imposed functional constraints under the Laplacian operator. With the properties of graph Laplacian, the nodal potentials *ϕ* can be solved using the Moore-Penrose pseudoinverse

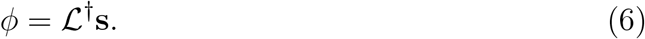

The eigendecomposition of the Laplacian ℒ = *U* Λ*U*^⊤^ where *U* = [**u**_1_, …, **u**_|*V* |_] is an orthonormal matrix whose columns **u**_*k*_ ∈ ℝ ^|*V* |^ are the eigenvectors of the graph Laplacian and Λ = diag(*λ*_1_, …, *λ*_|*V* |_) is the diagonal matrix of Laplacian eigenvalues leads to expression

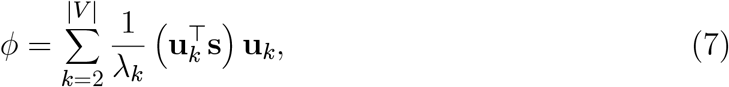

demonstrating that nodal potentials are linear combinations of Laplacian eigenmodes, weighted by the inverse of their eigenvalues. Indexing starts from *k* = 2 because the first eigenvalue of the graph Laplacian is zero corresponding to the constant mode which is excluded by the definition of the Moore-Penrose pseudoinverse. This spectral representation is related to prior work showing that low-frequency Laplacian eigenmodes dominate macroscale functional organization and capture indirect, polysynaptic interactions in the brain (Abdelnour et al., 2018, 2014; Atasoy et al., 2016; Deslauriers-Gauthier et al., 2020).

### 2.3 Mapping nodal information to edges with graph gradients

If the nodal potentials *ϕ* describe the spatial distribution of functional signals, the graph gradient operator ∇_*G*_ of it represents how the functional interactions are accomodated along structural pathways:

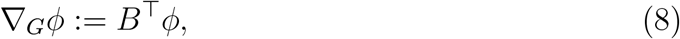

that assigns the difference in potential between incident nodes of each edge. Weighting this gradient by the edge-weight matrix *C* yields a definition for edge-wise flow **f** :

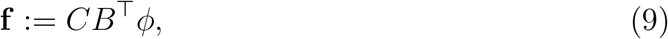

whose elements **f**_*e*_ = *e*_*w*_(*e*)(*ϕ*_*i*_ − *ϕ*_*j*_) quantifies how strongly a functional potential difference is supported by the corresponding structural connection. In this sense, edge weights act as pathway-specific gains that modulate the distribution of support under constraints.

Substituting the eigenmode expansion in eq. 7 into eq. 9 gives

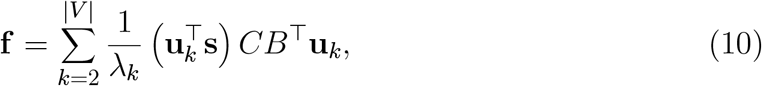

showing that edge flows correspond to weighted spatial derivatives of Laplacian eigenmodes, providing an explicit pathway-level realization of connectome harmonics. Therefore eq. 10 maps nodal functional information **s** onto structural graph *G*, yielding an edge-level representation consistent with the brain anatomy.

While existing structure–function coupling models implicitly account for multi-edge pathways, they typically summarize interactions at the node or region level, leaving the contributions of individual structural connections hidden (Greaves et al., 2025; Sporns, 2013; Suárez et al., 2020). To my knowledge, this edge-resolved spectral representation has not been previously explored in network neuroscience and therefore could provide new insights to brain network analyses.

The edge-wise flows defined in eq. 9 should be interpreted as a structural allocation of support required to satisfy the imposed pattern of pairwise functional relationships under the Laplacian constraint. In the present work, these relationships are derived from fMRI correlations, which reflect statistical dependencies rather than direct communication or causal interaction. Accordingly, a large flow on a given edge does not imply signal transmission but indicates that the edge lies on structurally efficient pathways that are consistent with the imposed constraints. More generally, the framework maps externally specified pairwise relationships onto the structural network, and the interpretation of the resulting edge-level quantities depends on the nature of the input measure. The practical interpretation of these quantities is elaborated further in the Discussion.

## 3 Methods

This section is organized as follows. First, the numerical framework underlying the proposed function-structure coupling is described 3.1. Next, a single-subject example is presented to illustrate tractogram filtering in the Default Mode Network (Raichle et al., 2001) 3.2. The framework is then evaluated using a combination of controlled and empirical analyses, including a structural specificity assessment using the FiberCup phantom 3.3, an in-silico function-structure simulation 3.4, a group-level analysis of 207 Human Connectome Project subjects (Van Essen et al., 2012) 3.5, an analysis of cross-subject variability 3.6, and a test-retest reproducibility analysis 3.7.

### 3.1 Constrained Laplacian as Modified Nodal Analysis problem

The theoretical formulation in Section 2 formulated function-structure coupling as a constrained Laplacian problem ℒ*ϕ* = **s**, where the unknowns are the nodal potentials *ϕ*, and **s** encodes externally imposed constraints. In practice, these constraints originate from selected functional connectivity (FC) values. The purpose of this section is to describe how these constraints are incorporated numerically using Modified Nodal Analysis (MNA) (Ho et al., 1975; Wedepohl and Jackson, 2002) in a manner consistent with the Laplacian formulation.

#### 3.1.1 Mapping functional connectivity to constrain values

Functional connectivity provides externally imposed pairwise constraints on differences between nodal potentials. The encoding of these constraints can be done with the following logic. An incidence matrix **B** ∈ ℝ^*n×m*^, whose columns specify which node pairs are constrained, and a vector of constraint values *s*_fc_ ∈ ℝ^*m*^, where each entry corresponds to the functional connectivity value assigned to that pair.

The circuit analogy used in this work is intended as a mathematical formalism rather than a biophysical model. Functional connectivity values serve as externally specified pairwise constraints, and the resulting potential- and current-like quantities represent how the structural network accommodates these constraints under the Laplacian’s conservation laws. No physical interpretation of these quantities is implied; the formulation draws on circuit theory only as a principled computational framework for expressing constraints and deriving edge-level representations.

In all experiments, FC values were Fisher z-transformed and used directly as imposed potential differences. All constraints are solved jointly within a single system, meaning no pairwise estimation or iterative inference is performed. For clarity, the theoretical nodal source term in Eq.5 is named as *s* whereas *s*_fc_ is the empirical constraint vector derived from functional connectivity.

#### 3.1.2 Modified Nodal Analysis (MNA) formulation

Rather than solving the constrained system via explicit pseudoinversion or eigendecomposition, MNA is used as it results in an equivalent but computationally efficient sparse linear system and physically interpretable results. With the structural Laplacian ℒ, the constraint incidence matrix **B**, and constraint values *s*_fc_, the forward problem is written as:

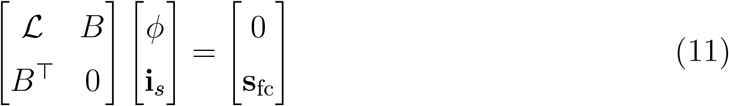

where *ϕ* ∈ ℝ^*n*^ are unknown nodal potentials (same as in Section 2), **i**_*s*_ ∈ ℝ^*m*^ are auxiliary variables that enforce the constraints (Ho et al., 1975; Wedepohl and Jackson, 2002), and the block *Bi*_*s*_ in the top equation plays the role of the effective nodal source term enforcing the imposed functional relationships. To avoid singularity, one node is grounded (e.g., *ϕ*_1_ = 0) This system is symmetric, semi-definite, and sparse, ensuring computational efficiency even for large connectomes.

This formulation is mathematically equivalent to solving ℒ*ϕ* = **s**, but avoids explicit pseudoinversion and allows functional constraints to be expressed directly as potential differences without redefining the Laplacian.

#### 3.1.3 Graph edge flows

Once nodal potentials *ϕ* are solved using MNA, flows along structural edges can be computed using eq. 9 or in element-wise form:

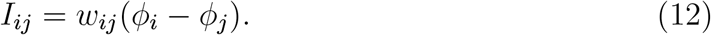

These values define an edge-centric connectivity matrix *I* that quantifies how support for the imposed functional constraints is distributed across structural connections. In the absence of functional constraints, the formulation reduces to a homogeneous Laplacian system equivalent to standard diffusion models of spectral graph theory. These flows represent how the structural network supports the externally imposed functional constraints. Importantly, the model does not attempt to infer directed influences or propagate activity; it simply computes the flow field implied by the constrained Laplacian.

### 3.2 Default Mode Network example

To demonstrate the proposed function-structure coupling method, one subject was selected from the Human Connectome Project (HCP) because this dataset provides both preprocessed resting-state fMRI (including ICA-derived components) and diffusion-weighted MRI tractography. This combination enables direct comparison of functional and structural connectivity within the same individual.

The Default Mode Network (DMN) was chosen as a test case due to its well-characterized role in intrinsic brain activity and strong function-structure associations. A DMN atlas (Smith and Nichols, 2009) was used to identify the ICA component with the highest spatial correlation to the DMN template. The resulting mask was segmented into six regions of interest (ROIs): medial prefrontal cortex (mPFC), posterior cingulate cortex (PCC), left and right inferior parietal lobules (IPL), and left and right temporal lobes (TL).

Functional connectivity between these ROIs was computed as Pearson’s correlation coefficient of the average fMRI time series within each region. This produced a 6×6 functional connectivity matrix representing pairwise correlations.

Whole-brain tractography was performed using MRtrix (Tournier et al., 2019) with multi-shell, multi-tissue constrained spherical deconvolution (Dhollander et al., 2016; Smith et al., 2012). Parameters included one million streamlines, minimum and maximum lengths of 5 mm and 250 mm, and a normalized white matter FOD cutoff of 0.05. The structural connectivity matrix was constructed by counting streamlines connecting each pair of DMN ROIs.

The functional and structural matrices were coupled using the MNA-based approach described in Section 3.1. Nodal voltages were solved from the simplified MNA system (eq. 11), and currents along structural edges were computed using eq. 12. These currents represent the estimation how the of functional information is accommodated across anatomical pathways.

The resulting current matrix highlights which structural connections receive strongest support under the functional constraints. For visualization, tractograms were filtered to retain streamlines with non-zero currents, and streamline-wise currents are shown with color scaling. This provides an intuitive representation of how functional relationships are supported across the structural network.

This single-subject example illustrates the feasibility of the proposed method and sets the stage for group-level analyses described in Section 2.4.

### 3.3 Structural specificity

To assess whether the proposed framework correctly maps imposed pairwise relationships onto anatomically appropriate structural pathways, a controlled validation was performed using the FiberCup diffusion MRI phantom (Fillard et al., 2011; Poupon et al., 2010, 2008). This dataset provides a known ground-truth fiber architecture, enabling evaluation under conditions where the underlying structural connectivity is explicitly defined.

Diffusion-weighted data from the phantom were processed using MRtrix3 (Tournier et al., 2019) with constrained spherical deconvolution to generate a whole-phantom tractogram containing one million streamlines. This tractogram was used to construct a structural connectivity matrix between the 12 regions shown as spherical nodes in Figure 4B.

Pairwise functional relationships were externally specified on a subset of node pairs defining five subnetworks (Figure 4A). Four pairs correspond to directly connected regions (subnetworks 1-4), while the fifth represents an indirect relationship between two regions without a direct structural connection, requiring a pathway through an intermediate node (node 5.3). A U-shaped bundle forms one of the subnetwork 4 is structurally isolated from the rest of the phantom and serves as a control: support should remain confined to the bundle and not propagate into unrelated pathways. All other entries of the constraint matrix were set to zero, and equal values were assigned to all constrained pairs to isolate the structural response of the model.

Given these imposed relationships and the structural connectivity matrix, the proposed framework was applied to compute nodal potentials and edge-level flows. These flows were used to filter the tractogram by retaining only streamlines consistent with the imposed relationships, allowing assessment of whether support is correctly allocated to direct bundles, routed through appropriate intermediate pathways in indirect cases, and absent from anatomically unrelated structures.

This experiment evaluates structural specificity, defined as the ability of the method to recover anatomically correct pathways given a known set of pairwise relationships. It does not constitute a full validation of the function-structure framework, as the imposed relationships do not arise from measured functional data, but instead provides a controlled baseline against which more realistic validation settings can be compared.

### 3.4 In-silico function-structure simulation

The in-silico simulation and the structural specificty experiment of the section 3.3 serve complementary validation roles and differ in how functional relationships are defined. In the section 3.3, pairwise relationships are externally specified and directly imposed, enabling a controlled test of whether the framework maps them onto the correct anatomical pathways. In contrast, the in-silico simulation generates functional connectivity from stochastic dynamics on the same structural scaffold, where activity propagates through the network to produce correlations between regions. Additional coupling is introduced for the same node pairs (Figure 4) to embed target relationships within the dynamics. Thus, the phantom isolates structural mapping properties, whereas the in-silico simulation evaluates the framework in a function-structure setting closer to empirical data.

The structural connectivity matrix derived from the phantom tractography in the section 3.3 was used as the coupling matrix of a linear dynamical system. To ensure stability, it was normalized by its spectral radius and scaled by a global coupling parameter (*α* = 0.1). Additional coupling was imposed on selected node pairs corresponding to the same subnetworks as in the phantom experiment, creating a controlled test case where success and failure are clearly defined. The coupling strength *β* was set to the 95th percentile of the nonzero entries of the structural matrix, ensuring detectability while remaining consistent with the scale of the underlying connectivity.

Node-wise time series were generated using the discrete-time system *x*(*t* + 1) = *Ax*(*t*) + *ϵ*(*t*), where *ϵ*(*t*) is Gaussian noise (Galán, 2008). Simulations were run for 5000 time steps, with the first 800 discarded as burn-in. The noise amplitude (*σ* = 0.01) was chosen to provide weak stochastic input relative to coupling strength, preserving structured interactions while avoiding deterministic behavior.

To obtain BOLD-like signals, the simulated time series were convolved with a canonical hemodynamic response function (HRF) following the standard SPM formulation (Friston et al., 1998), and sampled at TR = 0.72 s. Functional connectivity was computed as Pearson correlation between node time series, followed by Fisher z-transformation, yielding a matrix comparable to the empirical data.

The resulting structural and functional connectivity matrices were used as inputs to the proposed framework to compute nodal potentials and edge-level flows. This setup provides a complementary validation in which functional relationships arise from structure-driven dynamics, allowing assessment of whether the framework recovers the expected direct and indirect pathways under more realistic conditions.

### 3.5 Group study example

To assess the generalizability of the proposed method, it was applied to a cohort of 207 unrelated healthy adults from the HCP (Larivière et al., 2020; Van Essen et al., 2012). This group-level analysis demonstrates how the new dimension from the function-structure coupling produces a complementary connectivity matrix at the population scale.

Functional and structural connectivity matrices were obtained using the ENIGMA Toolbox (Larivière et al., 2021). The Desikan-Killiany atlas was selected for parcellation, resulting in 68 cortical regions per subject. Functional connectivity was computed from resting-state fMRI as Fisher z-transformed correlations, while structural connectivity was derived from diffusion MRI tractography.

Subject-level matrices were averaged to create group-level structural and functional connectomes. These were then coupled using the MNA-based approach described in Section 3.1 to compute the new connectome from eq. 12.

### 3.6 Cross-subject variability analysis

To assess subject-specific variability in the derived edge flows, an additional analysis was performed on the three HCP subjects (130013, 106824, and 419239) with the same preprocessing and tractography steps used for the DMN example. For each subject, the FSC model was applied separately to obtain an edge-flow matrix. To enable pattern-level comparisons across subjects, each matrix was vectorized (upper triangle) and z-scored per subject. Between-subject similarity was quantified using Pearson correlation of the vectorized flow patterns, overlap of the top 5% highest-flow edges using the Jaccard index, and edge-wise variance mapped back to matrix form. These metrics were computed analogously for structural and functional connectomes to contextualize flow variability relative to underlying SC and FC differences.

### 3.7 Test-retest reliability

To assess within-subject reproducibility, repeated resting-state acquisitions from the same three Human Connectome Project subjects described in Section 3.6 were analyzed, using identical structural connectivity and preprocessing. For each subject, four functional connectivity matrices (1LR, 1RL, 2LR, 2RL) were processed separately and used as inputs to the FSC framework, yielding corresponding edge-flow matrices.

Reproducibility was evaluated by comparing edge-level patterns across runs within subject. For each matrix, the upper triangle was vectorized, and similarity between runs was quantified using Pearson correlation and the Jaccard index applied to the top 5% of edges. Pairwise similarity matrices were computed for visualization.

## 4 Results

The proposed framework was evaluated using both empirical and controlled datasets. Human Connectome Project (HCP) data were used to demonstrate in vivo applicability, including a cohort of 207 subjects for group-level analysis, and a three-subject analysis of cross-subject variability and test-retest analysis in FSC-derived edge-level patterns (Larivière et al., 2021; Van Essen et al., 2012). The FiberCup diffusion phantom (Fillard et al., 2011; Poupon et al., 2010, 2008) was used to assess structural specificity under known ground-truth conditions, and an in-silico simulation on the same structural scaffold was performed to evaluate the method when functional connectivity arises from simulated dynamics.

### 4.1 Default Mode Network example

To illustrate the practical application of the proposed function-structure coupling framework, the Default Mode Network (DMN) from a HCP subject (100307) was analyzed. This dataset was selected because it provides both diffusion MRI tractography and preprocessed resting-state fMRI, enabling direct comparison of structural and functional connectivity within the same individual.

The DMN was chosen as a test case due to its well-established role in intrinsic brain activity and its strong function-structure associations. An ICA volume with the highest spatial correlation to the DMN atlas (Smith and Nichols, 2009) was identified, spatially smoothed, and thresholded to produce subject-specific DMN nodes, shown in figure 1. The six regions identified correspond approximately to the medial prefrontal cortex (mPFC), posterior cingulate cortex (PCC), bilateral inferior parietal lobules (IPL), and bilateral temporal lobes (TL). This experiment is not intended to redefine the DMN but to demonstrate how the proposed coupling algorithm can be applied to any network configuration.

**Figure 1:**
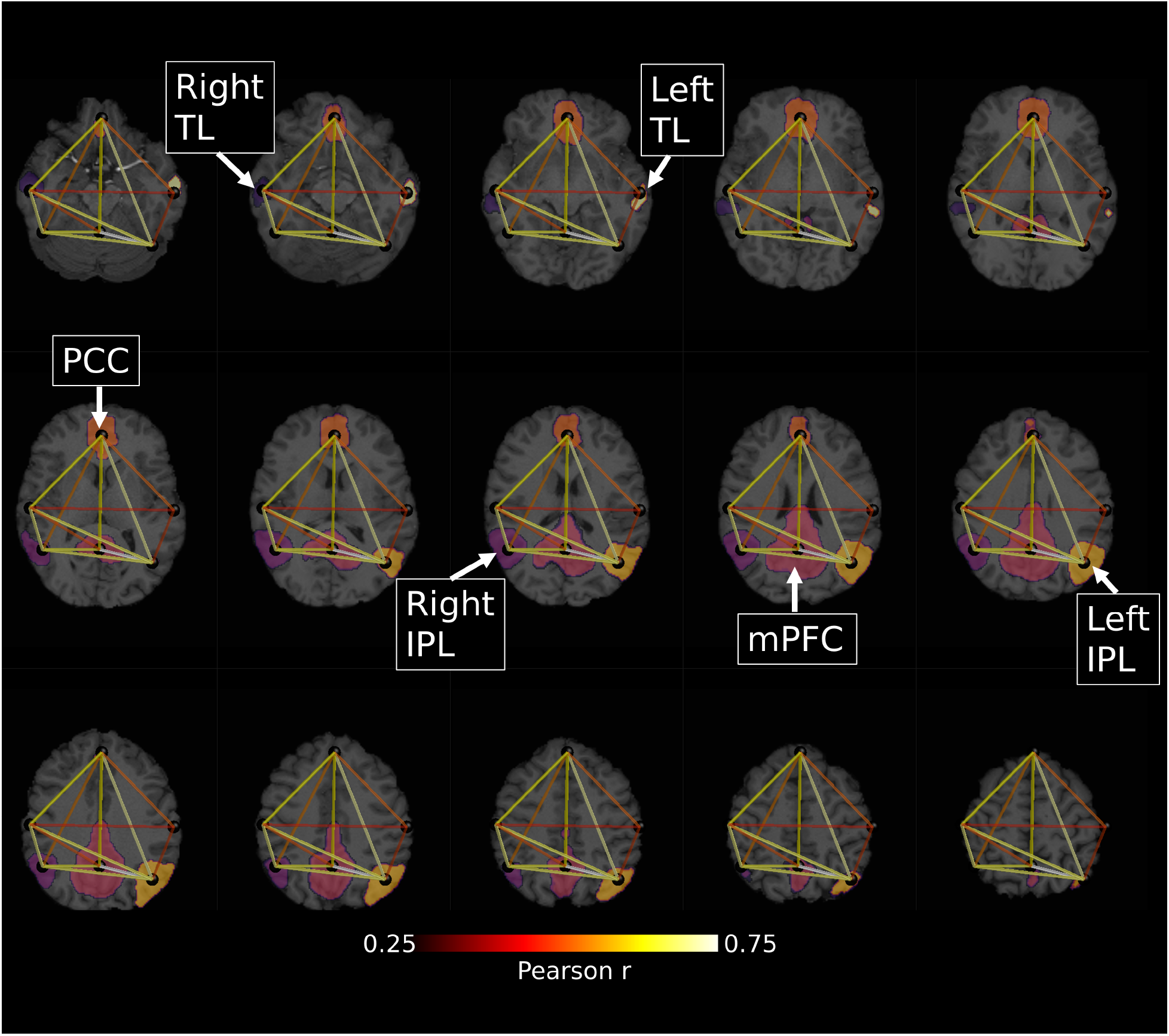
Visualization of the subject’s Default Mode Network (DMN). Six regions of interest are shown in distinct colors: medial prefrontal cortex (mPFC), posterior cingulate cortex (PCC), left and right inferior parietal lobules (IPL), and left and right temporal lobes (TL). The accompanying graph depicts functional connectivity between these regions, with edge color indicating the strength of functional correlations.

The subject’s DMN mask was used to extract all diffusion MRI tractography streamlines connected to the DMN, as shown in figure 3A. These streamlines were then filtered using the function-structure coupling approach, removing those with zero or near-zero current, as illustrated in figure 2B. Streamline-wise currents were applied as a colormap in the latter image to highlight which structural pathways support the imposed functional relationships. Finally, Figure 3 illustrates how streamline-wise currents can be visualized alongside nodal potentials, yielding an interpretable representation of how functional coupling is distributed across anatomical connections.

**Figure 2:**
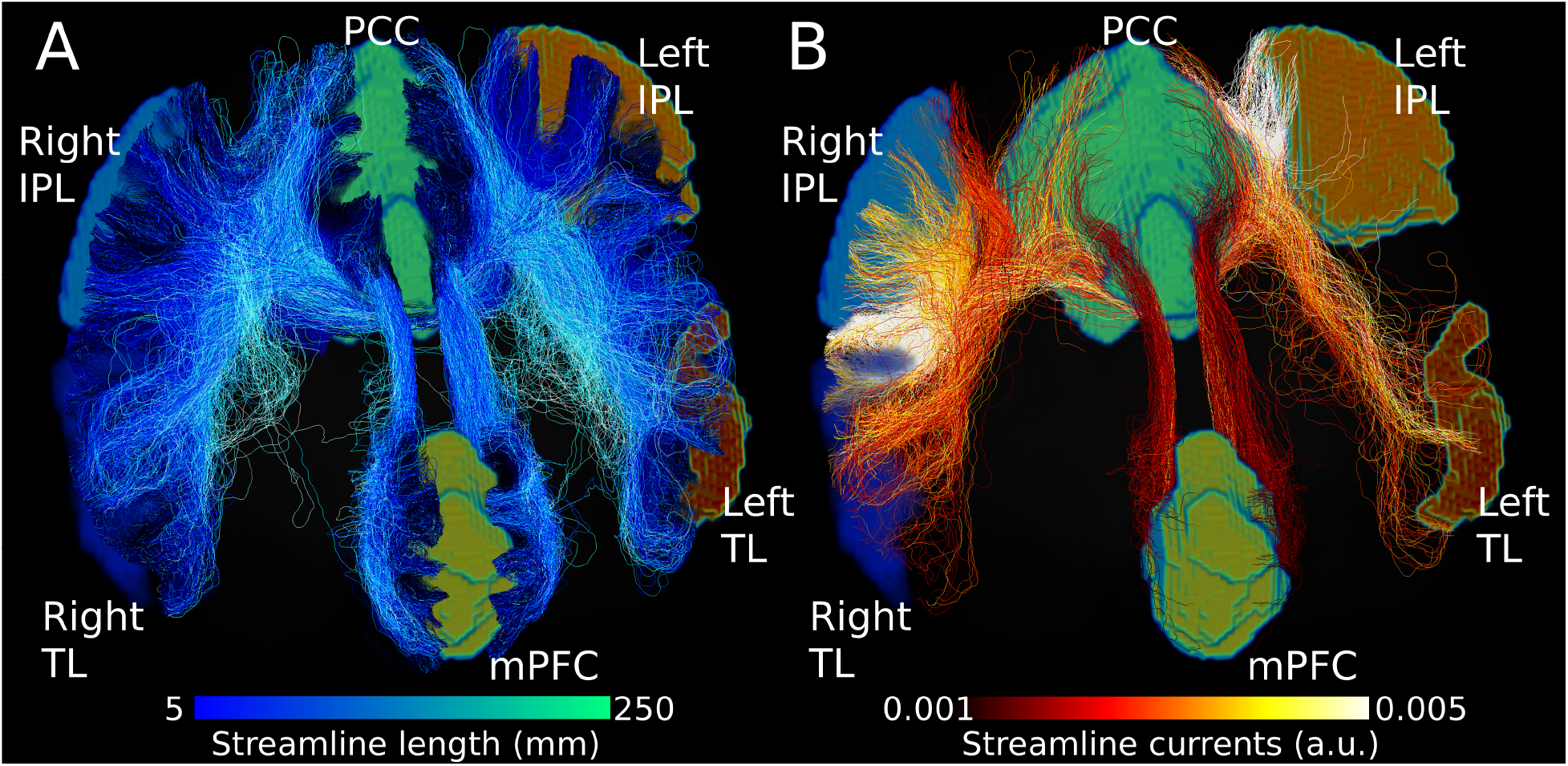
Tractography filtering using constrained Laplacian structure–function coupling in a single subject Default Mode Network (DMN). (A) Original tractogram connecting the six DMN regions, with streamline color indicating length. (B) Tractogram after filtering based on the current-based coupling, where streamlines with negligible current have been removed. Remaining streamlines are color-coded by streamline-wise current magnitude. The reduction in streamlines reflects selective retention of structural pathways that efficiently support the observed functional relationships, rather than uniform use of all anatomically reconstructed connections. This sparsification emphasizes pathways most consistent with the functional coupling pattern while suppressing weak or redundant trajectories inherent to tractography.

**Figure 3:**
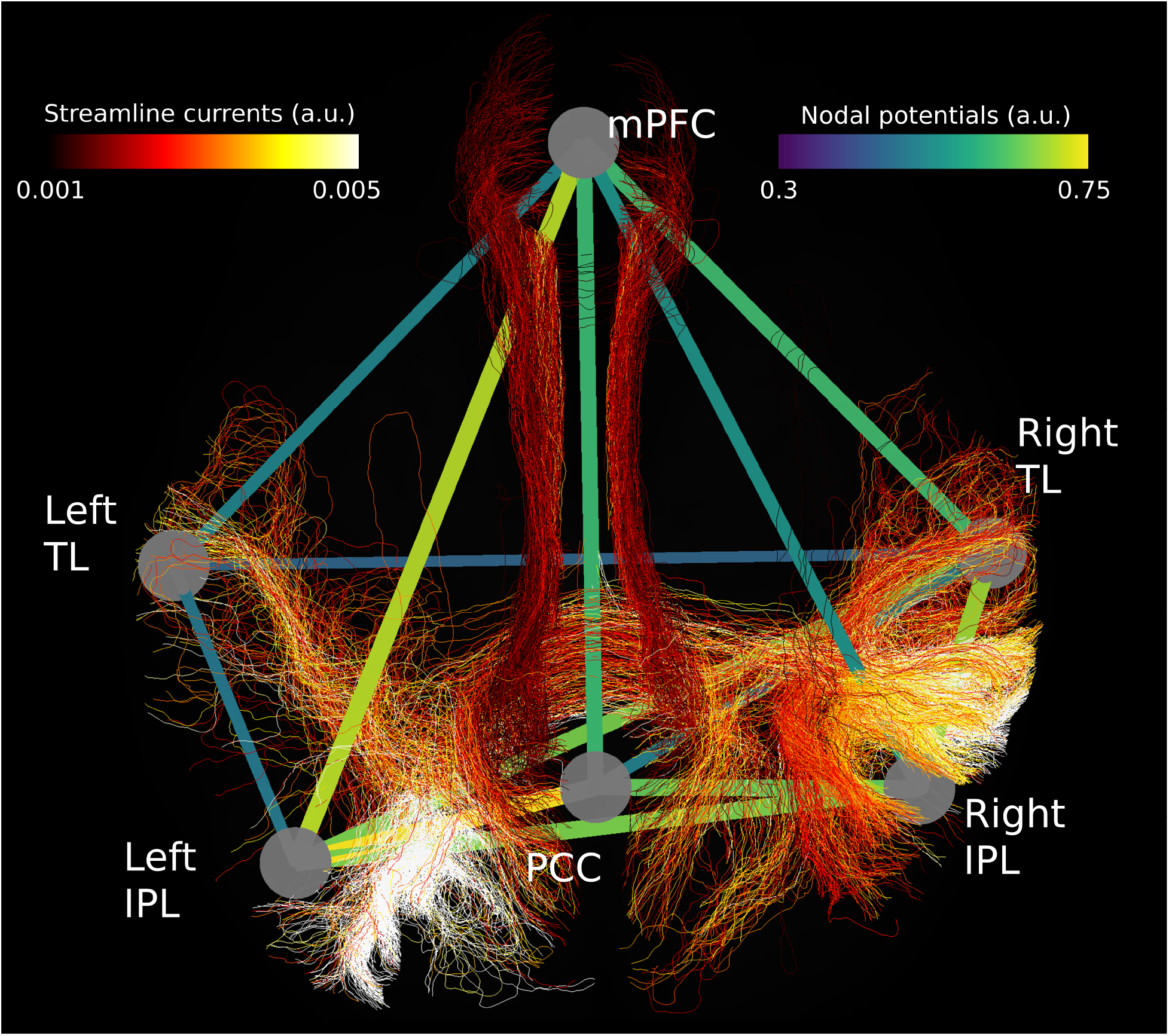
Visualization of nodal voltages and streamline-wise currents within the subject’s DMN. The graph depicts nodal voltages, with edges representing voltage drops between regions. Overlaid tractography streamlines are color-coded by corresponding functional current values, highlighting structural pathways that receive the strongest support. This combined representation provides an interpretable view of how functional relationships are supported by anatomical connections.

**Figure 4:**
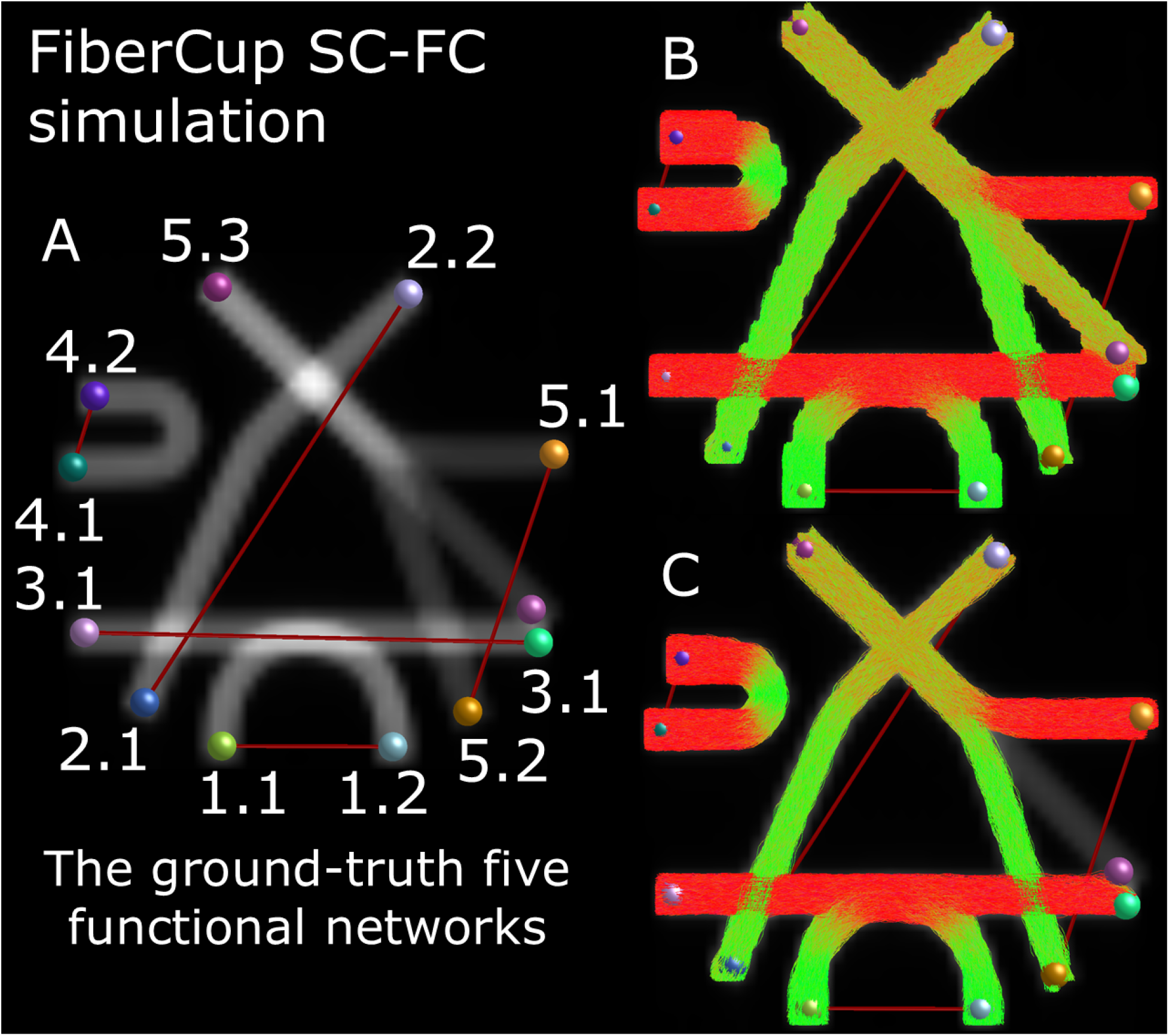
Ground-truth structural scaffold and resulting edge-level support patterns in the FiberCup dataset. Panel A illustrates the phantom divided into five predefined subnetworks, including an indirect relationship requiring a pathway from 5.1 to 5.2 through an intermediate node 5.3. Panel B shows the full tractogram derived from diffusion MRI, representing the complete structural connectivity. Panel C shows the filtered tractogram obtained using the proposed function-structure coupling, where only streamlines consistent with the imposed relationships are retained. The same qualitative behavior was observed in both the controlled phantom experiment (with externally specified relationships) and the in-silico simulation (where functional relationships emerge from simulated dynamics). In both cases, the framework correctly identifies direct structural pathways and routes indirect relationships through appropriate intermediate regions while avoiding anatomically unrelated structures.

### 4.2 Structural specificity

The controlled phantom experiment demonstrated that the proposed framework correctly maps imposed pairwise relationships onto anatomically appropriate structural pathways. For relationships corresponding to direct structural connections, support was concentrated on the known fiber bundles linking the associated regions. For the relationship imposed between regions without a direct connection, support was distributed along the indirect route via the appropriate intermediate node.

Application of the coupling to the tractogram resulted in the removal of structural segments not consistent with the imposed relationships, leaving only the relevant pathways. The U-shaped bundle, which forms an isolated structure with an imposed relationship between its endpoints, exhibited support confined to that bundle without spreading into unrelated pathways. This indicates that the method preserves anatomical specificity while avoiding spurious connections.

These results establish the structural specificity of the framework under controlled conditions. The same qualitative behavior was observed in the in-silico simulation described below, in which functional relationships arise from simulated dynamics rather than being externally specified.

### 4.3 In-silico function-structure simulation

When functional connectivity was generated from simulated node dynamics on the FiberCup structural scaffold, the proposed framework identified structural pathways consistent with all imposed relationships. Figure 4 shows the full tractogram and the corresponding filtered representation obtained using the coupling, in which streamlines not consistent with the imposed relationships are removed.

At the level of individual relationships (Figure 5), direct connections are associated with concentrated support on the corresponding fiber bundles, whereas relationships without a direct anatomical link are supported via indirect pathways through intermediate regions. In particular, the fifth case demonstrates that support is allocated along the correct indirect route, consistent with the known structural organization. Streamline-wise current magnitudes are lower for indirect pathways, reflecting their longer and less efficient structural routes.

**Figure 5:**
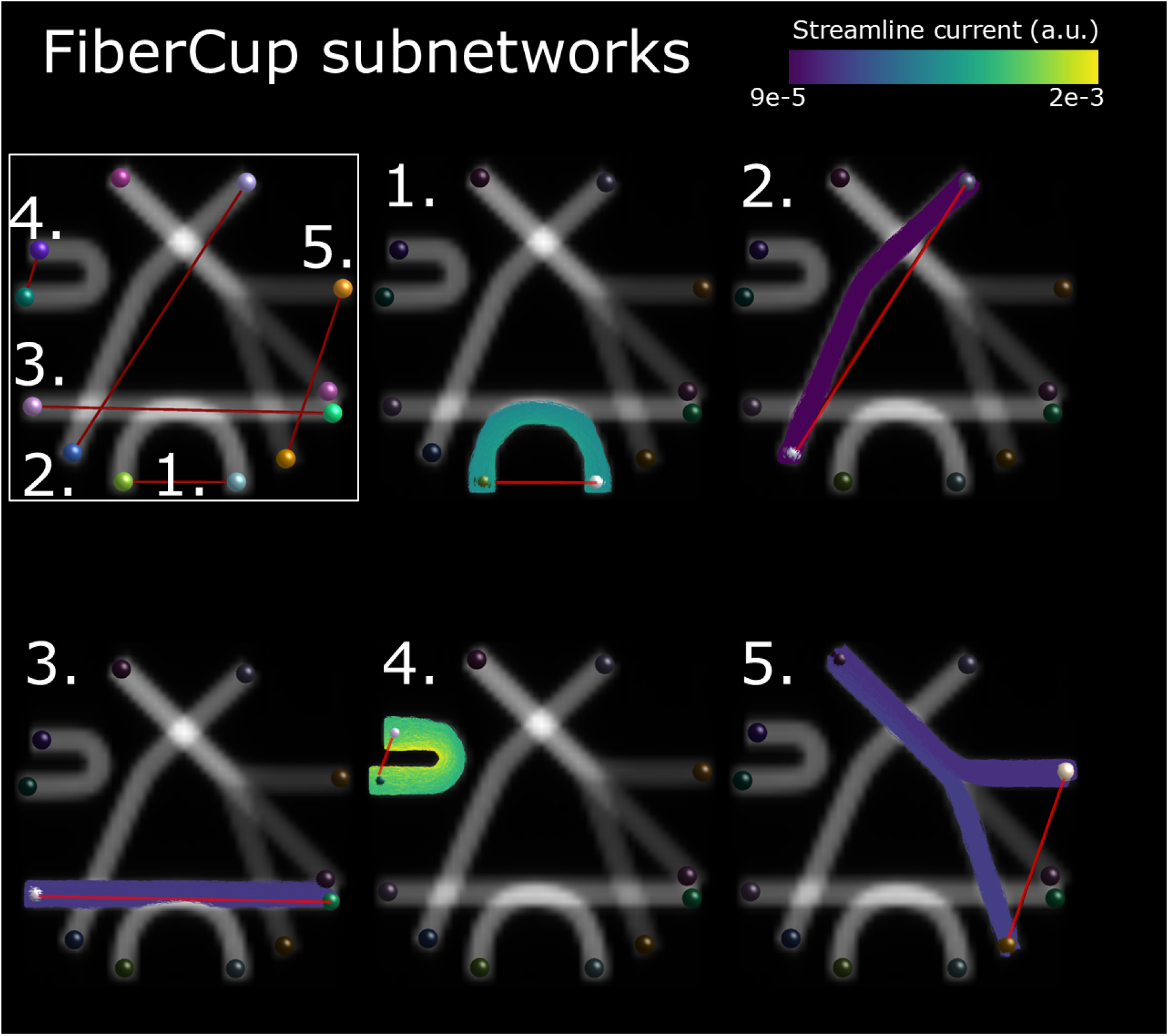
Subnetwork-level analysis of the FiberCup structural scaffold based on the in-silico simulation. Each panel shows the structural pathways receiving the strongest support for one of the five imposed pairwise relationships after applying the proposed coupling. All relationships are correctly associated with their corresponding anatomical pathways without spurious connections. The fifth case, which requires an indirect route from 5.1 to 5.2, is supported via the appropriate intermediate region 5.3. Color coding reflects streamline-wise current magnitude, with lower values observed for indirect pathways due to their longer and less efficient structural routes. The same qualitative behavior was observed in the controlled phantom experiment with externally specified relationships.

The qualitative behavior observed in this simulation matches that of the controlled phantom experiment. In both settings, the framework identifies direct structural pathways and distributes support across indirect routes when required, while avoiding anatomically unrelated structures. Because the resulting patterns were effectively identical, only the in-silico results are shown for brevity.

These results demonstrate that the framework produces consistent structural mappings not only for externally specified relationships, but also when functional connectivity arises from simulated dynamics on the structural network.

### 4.4 Group studies

To assess the generalizability of the proposed method, ít was applied to a cohort of 207 unrelated healthy adults from the Human Connectome Project (Van Essen et al., 2012). Group-level structural and functional connectivity matrices were computed using the Desikan–Killiany atlas and averaged across subjects (Larivière et al., 2021). These matrices were then coupled using the proposed framework to derive a current-based connectome.

Figure 6 compares the functional (red) and structural (blue) connectivity matrices with the resulting current-based connectome (green). Notably, the current-based connectome appears sparser than either of the original matrices, suggesting that functional relationships often converge along shared structural pathways rather than being distributed across all anatomical connections. This group-level analysis demonstrates that the proposed approach provides a complementary connectivity representation at the population scale, offering insights into how functional relationships are supported by structural networks.

**Figure 6:**
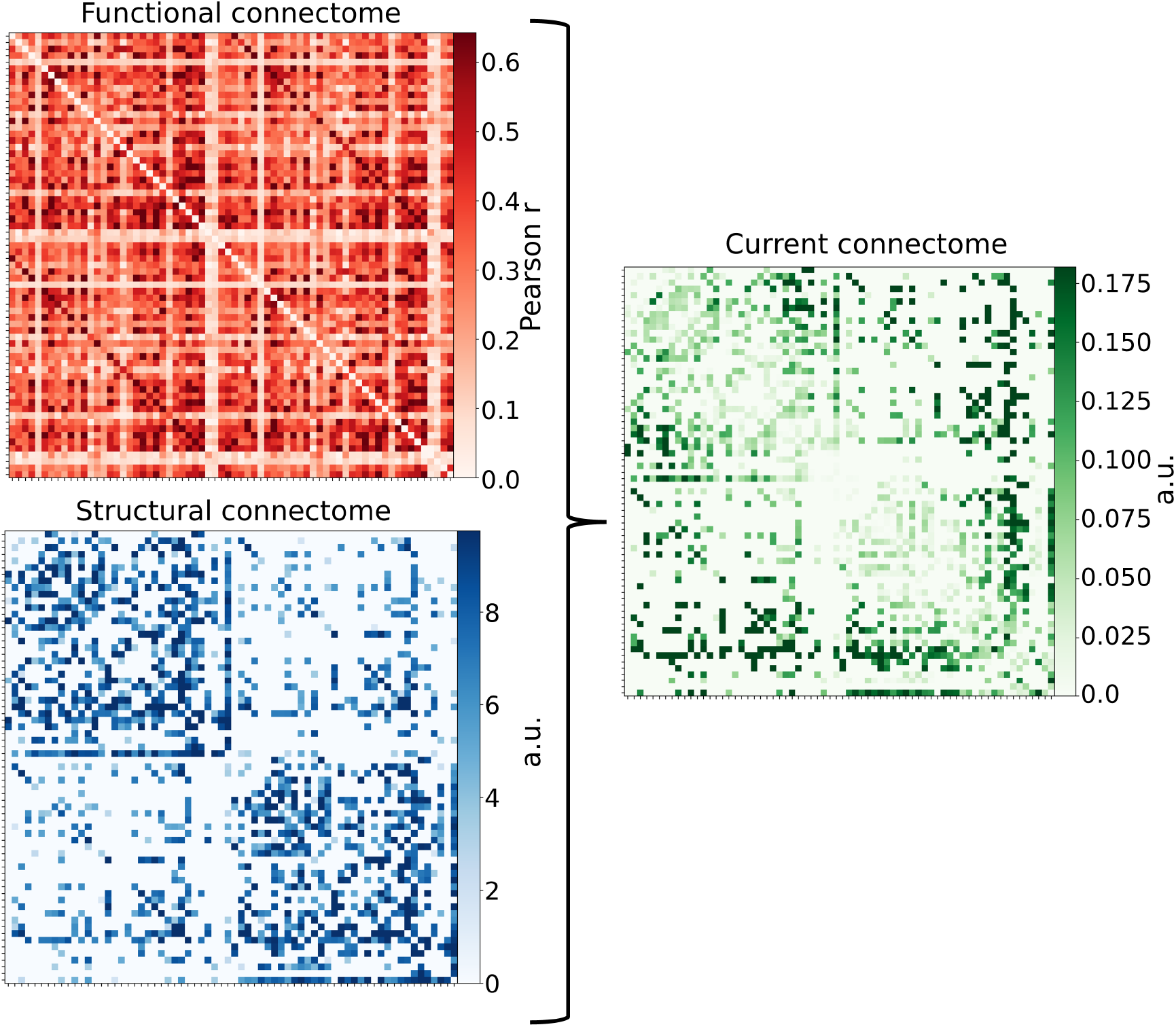
Group-level connectome analysis of 207 subjects from the Human Connectome Project. Functional connectivity (red) and structural connectivity (blue) matrices are shown alongside the resulting current-based connectome (green) derived from the proposed constrained Laplacian coupling. The current-based connectome is notably sparser than either the functional or structural matrices, reflecting that functional relationships are preferentially supported by a subset of anatomically efficient pathways rather than uniformly distributed across all available connections. This sparsity indicates convergence of functional interactions along shared structural routes, yielding a pathway-resolved representation that highlights structurally plausible substrates of functional coupling while suppressing weak or redundant connections. The axes correspond to the 68 cortical labels of Desikan-Killiany atlas which are omitted as the exact labels are not necessary for this visualization.

### 4.5 Cross-subject variability analysis

The three-subject analysis revealed clear subject-specific differences in the edge-flow patterns produced by the proposed model. Although flows showed moderate Pearson similarity across subjects (0.87-0.90), the overlap of the strongest pathways was only partial (Jaccard 0.44-0.48), and the edge-wise variance map exhibited spatially structured differences rather than uniform noise (Figure 7, top row). For comparison, structural connectomes were more similar across subjects (Pearson 0.94-0.95; Jaccard 0.64-0.73), while functional connectomes were less similar (Pearson 0.64-0.72; Jaccard 0.38-0.40) (Figure 7, middle and bottom rows). These findings indicate that FSC-derived flows are not trivially inherited from SC or FC individually but capture individualized function-structure coupling patterns.

**Figure 7:**
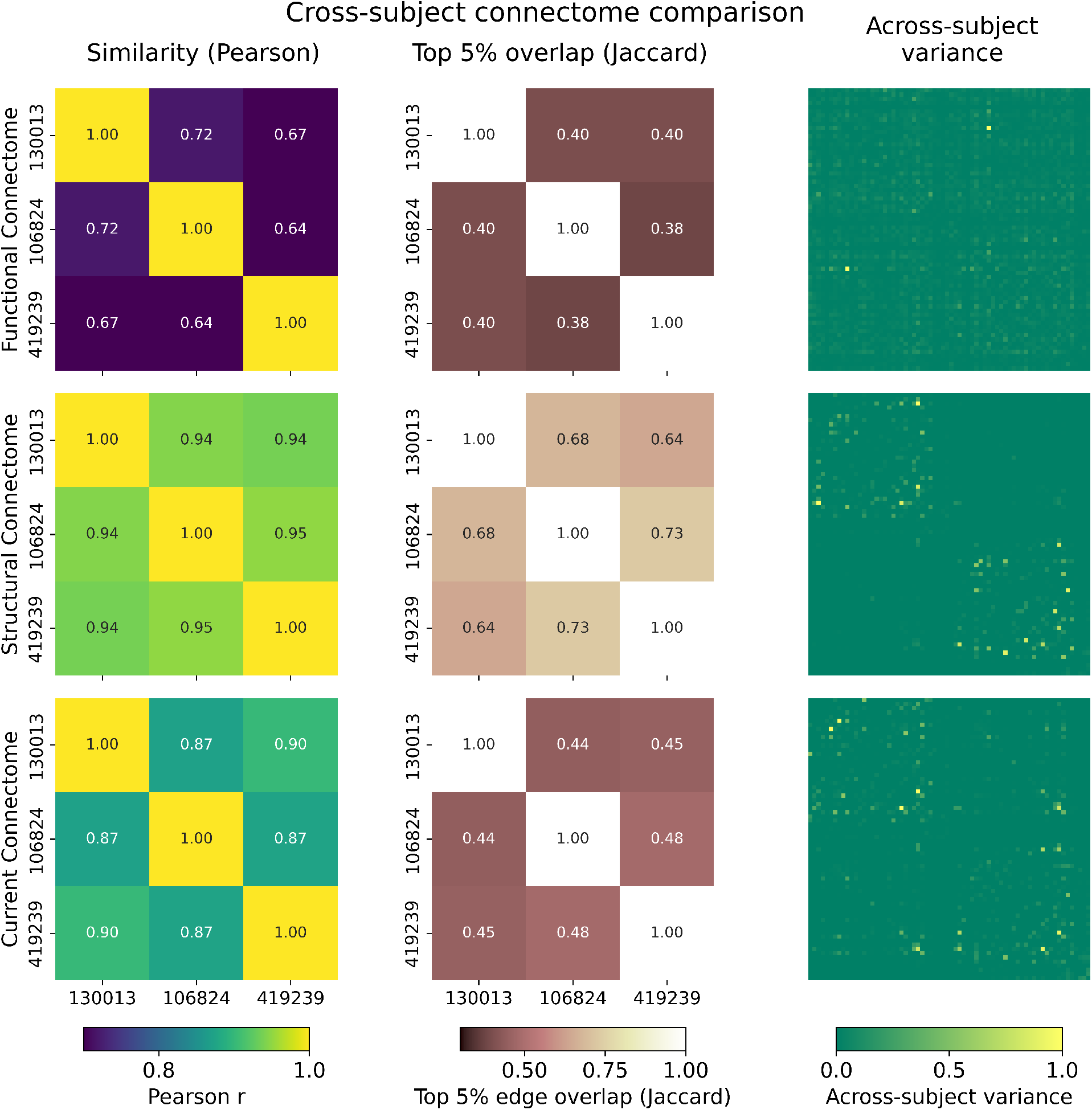
Each column summarizes between-subject similarity for three HCP participants (130013, 106824, 419239) using: (1) Pearson similarity of the vectorized matrices, (2) Jaccard overlap of the top-5% strongest edges, and (3) edge-wise variance across subjects where axes reprecent the 68 Desikan-Killiany labels. **Top row**: results for the proposed edge-flow formulation show moderate Pearson similarity (0.87-0.90) and only partial top-k overlap (Jaccard 0.44-0.48), together with spatially non-uniform variance, indicating clear subject-specific patterns in the distribution of functionally supported structural pathways. **Middle row**: structural connectomes exhibit higher cross-subject similarity (Pearson 0.94-0.95; Jaccard 0.64-0.73), reflecting shared anatomical organization. **Bottom row**: functional connectomes show lower similarity (Pearson 0.64-0.72; Jaccard 0.38-0.40), consistent with greater inter-subject variability in FC. Taken together, the comparison demonstrates that the proposed method produces individualized edge-flow signatures that are not trivially inherited from either SC or FC alone, but emerge from their joint interaction in the constrained Laplacian formulation.

### 4.6 Test-retest reliability

Within-subject reproducibility of functional connectivity and FSC-derived edge-flow patterns is shown in Figure 8. Across all subjects, functional connectivity exhibited moderate similarity between repeated runs, whereas the FSC-derived edge-flow matrices showed consistently higher agreement.

**Figure 8:**
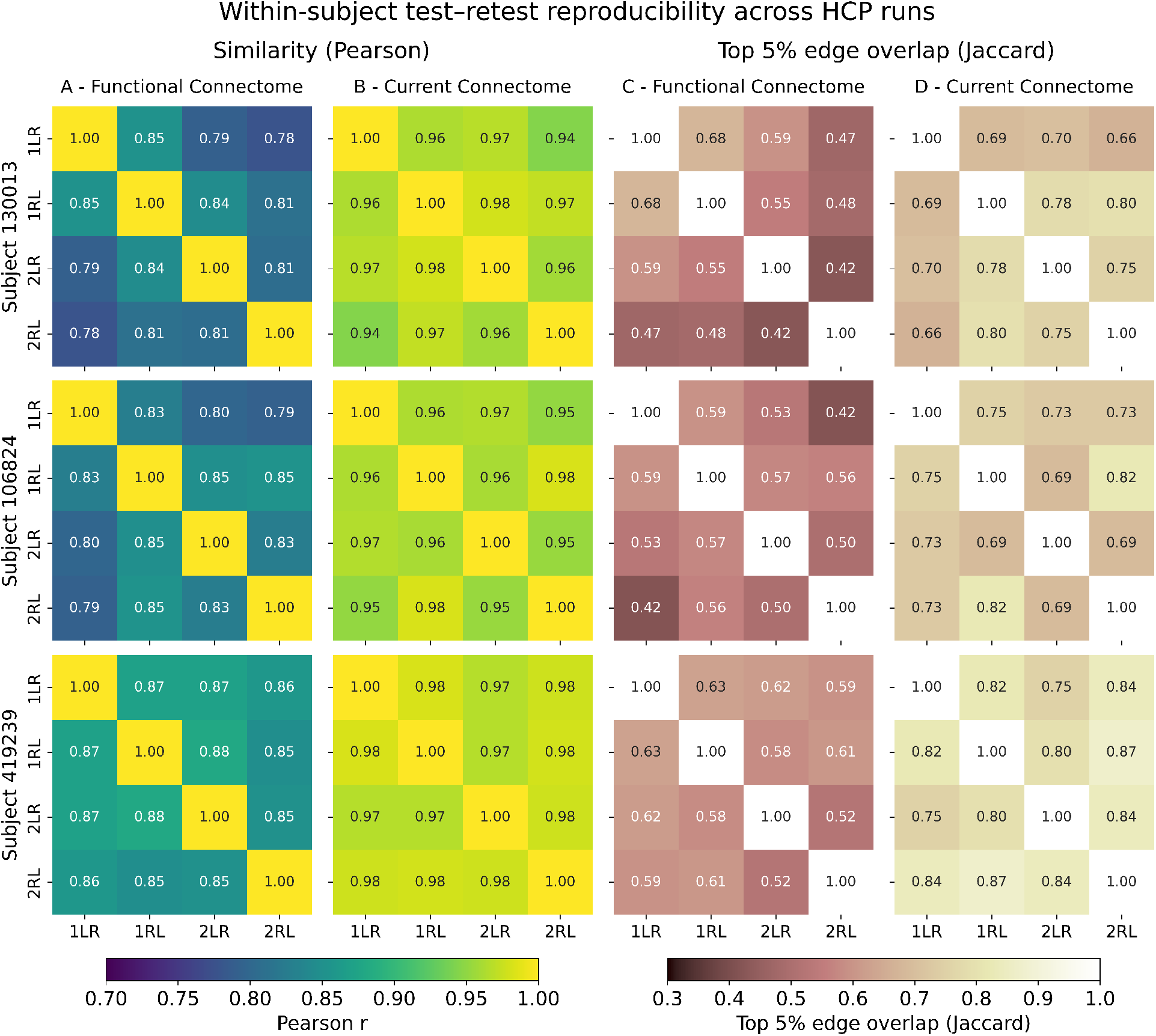
Within-subject test-retest reproducibility of functional connectivity (FC) and FSC-derived edge-flow patterns across four resting-state runs (1LR, 1RL, 2LR, 2RL) for three subjects. Columns A-B show Pearson correlation matrices for FC and FSC, and columns C-D show top 5% edge overlap (Jaccard index). FSC-derived patterns exhibit consistently higher similarity across runs than FC, both in overall edge-weight patterns and in the stability of the strongest connections.

Using Pearson correlation of vectorized upper-triangular entries, functional connectivity demonstrated correlations in the range of approximately 0.78-0.87 across runs, while the corresponding FSC-derived edge-flow patterns showed substantially higher correlations, typically exceeding 0.94 for all subjects and run pairs. A similar trend was observed for the top 5% edge overlap, where functional connectivity exhibited moderate Jaccard indices, whereas FSC-derived patterns showed consistently higher overlap across repeated runs.

This effect is visible in the full pairwise similarity matrices (Figure 8), where FSC-derived edge-flow patterns exhibit uniformly high similarity across all run combinations, in contrast to the more variable structure observed in functional connectivity. The increased reproducibility is observed both in overall edge-weight patterns (Pearson correlation) and in the stability of the strongest connections (Jaccard overlap).

These results indicate that the proposed framework produces edge-level representations that are more stable across repeated measurements than the underlying functional connectivity. This suggests that the integration of structural constraints with functional relationships reduces variability in the resulting edge-level patterns, while preserving the dominant connectivity structure observed across runs.

## 5 Discussion

This study investigated whether Modified Nodal Analysis (MNA), a method traditionally used in electrical circuit theory (Ho et al., 1975; Wedepohl and Jackson, 2002), can be adapted to study macroscale brain connectivity. The proposed framework is grounded in spectral graph theory (Chung, 1997; Shuman et al., 2013), reformulating the Laplacian constraint as an MNA system. This preserves flow conservation across nodes while extending the formulation to edge-level quantities, providing an interpretable representation of how structural pathways support imposed functional relationships. As with all function-structure approaches, the method inherits limitations from the underlying data, including tractography biases and fMRI noise.

Spectral graph theory has provided a principled foundation for modeling function-structure coupling, with approaches such as connectome harmonics and diffusion models leveraging Laplacian eigenmodes to capture indirect pathways and polysynaptic interactions (Abdelnour et al., 2018, 2014; Atasoy et al., 2016; Deslauriers-Gauthier et al., 2020). The proposed method retains this theoretical basis but introduces an edge-centric perspective, explicitly identifying which structural connections are most consistent with an observed pattern of functional relationships. This shift from node-level summaries to pathway-specific insights represents a conceptual advance, enabling tractography filtering and connectivity analyses that are both mathematically consistent and anatomically interpretable.

### 5.1 Interpretation of edge-level connectivity

The edge-level quantities (flows/currents) produced by the proposed framework should be interpreted as a structural allocation of support required to satisfy a pattern of pairwise functional relationships. This does not imply causal signal transmission, but rather identifies structurally plausible routes that are consistent with the observed statistical dependencies.

To make the interpretation of the model outputs concrete, consider a setting in which strong functional connectivity is observed between two regions, such as the posterior cingulate cortex (PCC) and medial prefrontal cortex (mPFC) in the default mode network. Applying the proposed method yields an edge-level representation indicating which structural connections are most strongly implicated in supporting this relationship.

This interpretation has potential practical implications. In a neurosurgical or brain stimulation context, this could be used to identify structural pathways whose disruption would be most consistent with altering the observed functional relationship. For instance, if a planned surgical resection intersects pathways assigned high support by the model, one may expect a greater impact on the associated functional coupling than if only low-support pathways are affected.

### 5.2 Validation and limitations

The validation strategy combines complementary analyses spanning controlled, simulated, and empirical settings. The FiberCup phantom provides a controlled test of structural specificity under known ground-truth conditions. The in-silico simulation extends this to a setting where functional connectivity emerges from dynamics on the structural network. Empirical validation includes a single-subject demonstration, group-level analysis across 207 Human Connectome Project subjects, cross-subject variability assessment, and test-retest analysis to evaluate reproducibility. Together, these experiments assess the framework’s behavior across idealized and realistic conditions, supporting its consistency and general applicability.

Across the controlled and simulated settings, the framework consistently assigns support to anatomically appropriate pathways for both direct and indirect relationships while avoiding spurious connections. This demonstrates that the method maps functional relationships onto structurally plausible routes, regardless of whether these relationships are externally imposed or arise from network dynamics.

The subject-specific variability analysis (Figure 7) indicates that the proposed framework captures individualized function-structure coupling rather than simply reproducing shared anatomical constraints. While SC displayed high similarity across subjects and FC showed substantial variability, the edge-flow patterns reflected a combination of both: moderately similar across individuals but with a meaningful proportion of subject-unique high-flow pathways. The spatial structure of the variance map further indicates that inter-subject differences arise in specific subnetworks rather than random fluctuations. This supports the suitability of FSC for individual-level mapping.

The test-retest analysis (Figure 8) further demonstrated that FSC-derived edge-level patterns are more reproducible than functional connectivity alone. Across repeated acquisitions, edge-flow representations exhibited consistently higher similarity, both in overall patterns and in the stability of the strongest connections. This suggests that incorporating structural constraints stabilizes the representation of interregional relationships, reducing variability while preserving dominant connectivity structure.

Diffusion MRI tractography inevitably contains false negatives and false positives, which raises questions about how function-structure models behave under incomplete or noisy anatomical input. The proposed constrained-Laplacian formulation is fully deterministic: it cannot generate structural connections that are not present. Missing tracts therefore remain missing, rather than being “filled in” by the model. An effect illustrated in the FiberCup phantom, where the isolated U-shaped bundle supports only its internally imposed relationship and does not receive support from other subnetworks. This reflects a core property of the method: it respects the topology of the supplied network rather than inferring new pathways.

False-positive streamlines also produce predictable behavior. Short spurious shortcuts attract disproportionately large flow and are identifiable as statistical outliers, whereas long or anatomically implausible false-positive tracts will be assigned with minimal flow and can be discounted accordingly. While exploring these effects systematically would require dedicated diffusion MRI simulation work beyond the scope of the present study, these characteristics clarify how the method behaves under typical tractography imperfections and how edge-level flow patterns can be interpreted in the presence of structural uncertainty.

The ability to compute streamline-wise currents (Figure 3) opens new opportunities for along-tract analyses (Colby et al., 2012), tractometry (Chamberland et al., 2021), and tractogram filtering (Sairanen et al., 2021). Identifying which white matter pathways support the functional relationships could provide insights into disorders that disrupt connectivity, such as aphasia, where structural changes vary across individuals (Sihvonen et al., 2024). This could support personalized rehabilitation strategies by pinpointing which functional routes are impaired or restored.

Importantly, the proposed framework produces an adjacency-like matrix, making it compatible with existing network analysis tools, including GNNs (Sporns, 2013). Similarly, tractography visualization platforms can readily display streamline-wise currents, as demonstrated in Figure 3. The open-source implementation provided with this study integrates seamlessly with these tools, facilitating adoption within the computational neuroscience community.

In the future, the framework could be extended into dynamic formulations by introducing time-dependent voltage sources and conductance changes, transforming the system in eq. 11 to complete MNA (Wedepohl and Jackson, 2002). This would align with dynamic causal modeling (Friston, 2011) and large-scale dynamic brain models (Breakspear, 2017; Deco, 2015), enabling simulation of transient neural activity and event-driven responses.

This would allow simulation of time series directly within the MNA framework, where time-varying activity at individual regions propagates through the structural network according to its topology. For example, transient inputs or perturbations applied at specific nodes could be used to study how activity spreads across anatomically constrained pathways and how edge-level currents evolve over time.

Such an extension would align the framework with dynamic causal modeling (Friston, 2011) and large-scale brain models (Breakspear, 2017; Deco, 2015), in which neural activity is modeled as a time-evolving process constrained by structural connectivity. By embedding these dynamics within the MNA formulation, the framework could provide a unified representation linking structural connectivity, time-resolved neural activity, and edge-level pathway contributions.

In summary, this work bridges spectral graph theory and circuit analysis to create a novel, interpretable representation of function-structure coupling. By preserving Laplacian-based principles while extending them to edge-level modeling, the proposed method provides both theoretical rigor and practical utility. Its interpretability offers unique advantages for tractography filtering and clinical applications. Future studies should explore dynamic extensions and validate the approach in patient populations to fully realize its potential.

## 6 Acknowledgements

V.S. was supported by the Orion Research Foundation sr and Instrumentarium Science Foundation sr. The authors wish to thank the Finnish Computing Competence Infrastructure (FCCI) for supporting this project with computational and data storage resources. Open access funded by Helsinki University Library.

## Footnotes

1 https://github.com/vilsaira/FSC

## Notes

### Competing Interest Statement

The authors have declared no competing interest.

### Summary of Updates

Added in-silico experiment and HCP test-retest analysis.

## References

Abdelnour, F., Dayan, M., Devinsky, O., Thesen, T., and Raj, A. (2018). Functional brain connectivity is predictable from anatomic network’s laplacian eigen-structure. NeuroImage, 172:728–739.

Abdelnour, F., Voss, H. U., and Raj, A. (2014). Network diffusion accurately models the relationship between structural and functional brain connectivity networks. Neuroimage, 90:335–347.

Atasoy, S., Donnelly, I., and Pearson, J. (2016). Human brain networks function in connectome-specific harmonic waves. Nature communications, 7(1):10340.

Breakspear, M. (2017). Dynamic models of large-scale brain activity. Nature neuroscience, 20(3):340–352.

Breakspear, M., Jirsa, V., and Deco, G. (2010). Computational models of the brain: from structure to function.

Calamante, F., Masterton, R. A., Tournier, J.-D., Smith, R. E., Willats, L., Raffelt, D., and Connelly, A. (2013). Track-weighted functional connectivity (tw-fc): a tool for characterizing the structural–functional connections in the brain. Neuroimage, 70:199–210.

Chamberland, M., Genc, S., Tax, C. M., Shastin, D., Koller, K., Raven, E. P., Cunningham, A., Doherty, J., van den Bree, M. B., Parker, G. D., et al. (2021). Detecting microstructural deviations in individuals with deep diffusion mri tractometry. Nature computational science, 1(9):598–606.

Chen, P., Yang, H., Zheng, X., Jia, H., Hao, J., Xu, X., Li, C., He, X., Chen, R., Okubo, T. S., et al. (2024). Group-common and individual-specific effects of structure–function coupling in human brain networks with graph neural networks. Imaging Neuroscience, 2:1–21.

Chung, F. R. (1997). Spectral graph theory, volume 92. American Mathematical Soc.

Colby, J. B., Soderberg, L., Lebel, C., Dinov, I. D., Thompson, P. M., and Sowell, E. R. (2012). Along-tract statistics allow for enhanced tractography analysis. Neuroimage, 59(4):3227–3242.

Deco, G. (2015). The dynamics of resting fluctuations in the brain. BMC Neuroscience, 16(Suppl 1):A2.

Deslauriers-Gauthier, S., Lina, J.-M., Butler, R., Whittingstall, K., Gilbert, G., Bernier, P.-M., Deriche, R., and Descoteaux, M. (2019). White matter information flow mapping from diffusion mri and eeg. NeuroImage, 201:116017.

Deslauriers-Gauthier, S., Zucchelli, M., Frigo, M., and Deriche, R. (2020). A unified framework for multimodal structure–function mapping based on eigenmodes. Medical Image Analysis, 66:101799.

Dhollander, T., Raffelt, D., and Connelly, A. (2016). Unsupervised 3-tissue response function estimation from single-shell or multi-shell diffusion mr data without a co-registered t1 image. In ISMRM workshop on breaking the barriers of diffusion MRI, volume 5. Lisbon.

Fillard, P., Descoteaux, M., Goh, A., Gouttard, S., Jeurissen, B., Malcolm, J., Ramirez-Manzanares, A., Reisert, M., Sakaie, K., Tensaouti, F., et al. (2011). Quantitative evaluation of 10 tractography algorithms on a realistic diffusion mr phantom. Neuroimage, 56(1):220–234.

Friston, K. J. (2011). Functional and effective connectivity: a review. Brain, 1(1):13–36. MAG ID: 2061564920.

Friston, K. J., Fletcher, P., Josephs, O., Holmes, A., Rugg, M. D., and Turner, R. (1998). Event-related fmri: characterizing differential responses. Neuroimage, 7(1):30–40.

Galán, R. F. (2008). On how network architecture determines the dominant patterns of spontaneous neural activity. PloS one, 3(5):e2148.

Greaves, M. D., Novelli, L., Mansour L. S., Zalesky, A., and Razi, A. (2025). Structurally informed models of directed brain connectivity. Nature Reviews Neuroscience, 26(1):23–41. Publisher: Nature Publishing Group.

Ho, C.-W., Ruehli, A., and Brennan, P. (1975). The modified nodal approach to network analysis. IEEE Transactions on Circuits and Systems, 22(6):504–509. Conference Name: IEEE Transactions on Circuits and Systems.

Honey, C. J., Sporns, O., Cammoun, L., Gigandet, X., Thiran, J.-P., Meuli, R., and Hagmann, P. (2009). Predicting human resting-state functional connectivity from structural connectivity. Proceedings of the National Academy of Sciences, 106(6):2035–2040.

Hong, Y., Cornea, E., Girault, J. B., Bagonis, M., Foster, M., Kim, S. H., Prieto, J. C., Chen, H., Gao, W., Styner, M. A., et al. (2023). Structural and functional connectome relationships in early childhood. Developmental Cognitive Neuroscience, 64:101314.

Huang, H. and Ding, M. (2016). Linking Functional Connectivity and Structural Connectivity Quantitatively: A Comparison of Methods. Brain, 6(2):99–108. MAG ID: 2229937548.

Larivière, S., Paquola, C., Park, B.-y., Royer, J., Wang, Y., Benkarim, O., Vos de Wael, R., Valk, S. L., Thomopoulos, S. I., Kirschner, M., Lewis, L. B., Evans, A. C., Sisodiya, S. M., McDonald, C. R., Thompson, P. M., and Bernhardt, B. C. (2021). The ENIGMA Toolbox: multiscale neural contextualization of multisite neuroimaging datasets. Nature Methods, 18(7):698–700. Number: 7 Publisher: Nature Publishing Group.

Larivière, S., Rodríguez-Cruces, R., Royer, J., Caligiuri, M. E., Gambardella, A., Concha, L., Keller, S. S., Cendes, F., Yasuda, C., Bonilha, L., Gleichgerrcht, E., Focke, N. K., Domin, M., von Podewills, F., Langner, S., Rummel, C., Wiest, R., Martin, P., Kotikalapudi, R., O’Brien, T. J., Sinclair, B., Vivash, L., Desmond, P. M., Alhusaini, S., Doherty, C. P., Cavalleri, G. L., Delanty, N., Kälviäinen, R., Jackson, G. D., Kowalczyk, M., Mascalchi, M., Semmelroch, M., Thomas, R. H., Soltanian-Zadeh, H., Davoodi-Bojd, E., Zhang, J., Lenge, M., Guerrini, R., Bartolini, E., Hamandi, K., Foley, S., Weber, B., Depondt, C., Absil, J., Carr, S. J. A., Abela, E., Richardson, M. P., Devinsky, O., Severino, M., Striano, P., Tortora, D., Hatton, S. N., Vos, S. B., Duncan, J. S., Whelan, C. D., Thompson, P. M., Sisodiya, S. M., Bernasconi, A., Labate, A., McDonald, C. R., Bernasconi, N., and Bernhardt, B. C. (2020). Network-based atrophy modeling in the common epilepsies: A worldwide ENIGMA study. Science Advances, 6(47):eabc6457. Publisher: American Association for the Advancement of Science.

Liu, Z.-Q., Vazquez-Rodriguez, B., Spreng, R. N., Bernhardt, B. C., Betzel, R. F., and Misic, B. (2022). Time-resolved structure-function coupling in brain networks. Communications biology, 5(1):532.

Neudorf, J., Kress, S., and Borowsky, R. (2022). Structure can predict function in the human brain: a graph neural network deep learning model of functional connectivity and centrality based on structural connectivity. Brain Structure and Function, 227(1):331–343.

Novelli, L., Friston, K., and Razi, A. (2024). Spectral dynamic causal modeling: A didactic introduction and its relationship with functional connectivity. Network Neuroscience, 8(1):178–202.

Poupon, C., Laribiere, L., Tournier, G., Bernard, J., Fournier, D., Fillard, P., Descoteaux, M., and Mangin, J.-F. (2010). A diffusion hardware phantom looking like a coronal brain slice. In Proceedings of the international society for magnetic resonance in medicine, volume 18, page 581.

Poupon, C., Rieul, B., Kezele, I., Perrin, M., Poupon, F., and Mangin, J.-F. (2008). New diffusion phantoms dedicated to the study and validation of high-angular-resolution diffusion imaging (hardi) models. Magnetic Resonance in Medicine: An Official Journal of the International Society for Magnetic Resonance in Medicine, 60(6):1276–1283.

Raichle, M. E., MacLeod, A. M., Snyder, A. Z., Powers, W. J., Gusnard, D. A., and Shulman, G. L. (2001). A default mode of brain function. Proceedings of the national academy of sciences, 98(2):676–682.

Sairanen, V., Ocampo-Pineda, M., Granziera, C., Schiavi, S., and Daducci, A. (2021). Incorporating outlier information into diffusion MR tractogram filtering for robust structural brain connectivity and microstructural analyses. Technical report. Company: Cold Spring Harbor Laboratory Distributor: Cold Spring Harbor Laboratory Label: Cold Spring Harbor Laboratory Section: New Results Type: article.

Shuman, D. I., Narang, S. K., Frossard, P., Ortega, A., and Vandergheynst, P. (2013). The emerging field of signal processing on graphs: Extending high-dimensional data analysis to networks and other irregular domains. IEEE signal processing magazine, 30(3):83–98.

Sihvonen, A. J., Pitkäniemi, A., Siponkoski, S.-T., Kuusela, L., Martínez-Molina, N., Laitinen, S., Särkämö, E.-R., Pekkola, J., Melkas, S., Schlaug, G., et al. (2024). Structural neuroplasticity effects of singing in chronic aphasia. Eneuro, 11(5).

Smith, R. E., Tournier, J.-D., Calamante, F., and Connelly, A. (2012). Anatomically-constrained tractography: improved diffusion mri streamlines tractography through effective use of anatomical information. Neuroimage, 62(3):1924–1938.

Smith, S. M. and Nichols, T. E. (2009). Threshold-free cluster enhancement: addressing problems of smoothing, threshold dependence and localisation in cluster inference. Neuroimage, 44(1):83–98.

Sporns, O. (2013). Structure and function of complex brain networks. Dialogues in Clinical Neuroscience, 15(3):247–262.

Suárez, L. E., Markello, R. D., Betzel, R. F., and Misic, B. (2020). Linking Structure and Function in Macroscale Brain Networks. Trends in Cognitive Sciences, 24(4):302– 315.

Tournier, J.-D., Smith, R., Raffelt, D., Tabbara, R., Dhollander, T., Pietsch, M., Christiaens, D., Jeurissen, B., Yeh, C.-H., and Connelly, A. (2019). Mrtrix3: A fast, flexible and open software framework for medical image processing and visualisation. Neuroimage, 202:116137.

Van Essen, D. C., Ugurbil, K., Auerbach, E. J., Behrens, T. E., Bucholz, R. D., R. Bucholz, Chang, A., Chen, L., Maurizio Corbetta, Corbetta, M., Della Penna, S., Curtiss, S. W., Della Penna, S., Feinberg, D. A., Glasser, M. F., Heath, A. C., Harel, N., Heath, A. C., Larson-Prior, L. J., Marcus, D. S., Michalareas, G., Moeller, S., Oostenveld, R., Petersen, S., Petersen, S. E., Prior, F. W., Schlaggar, B. L., Smith, S. M., Snyder, A. Z., Xu, J., and Yacoub, E. (2012). The Human Connectome Project: A data acquisition perspective. NeuroImage, 62(4):2222–2231. MAG ID: 2020519533.

Wedepohl, L. M. and Jackson, L. (2002). Modified nodal analysis: an essential addition to electrical circuit theory and analysis. Engineering Science & Education Journal, 11(3):84–92. Publisher: IET Digital Library.

Yang, H., Winter, S., Zhang, Z., and Dunson, D. (2022). Interpretable ai for relating brain structural and functional connectomes. arXiv preprint 2210.05672.

Zalesky, A., Sarwar, T., Tian, Y., Liu, Y., Yeo, B. T., and Ramamohanarao, K. (2024). Predicting an individual’s functional connectivity from their structural connectome: Evaluation of evidence, recommendations, and future prospects. Network Neuroscience, 8(4):1291–1309.

